# An improved gray whale assembly highlights how allospecific reference-genome choice can affect genomic diversity estimates

**DOI:** 10.1101/2025.01.23.634050

**Authors:** Anna Brüniche-Olsen, Andrew Black, Kimberly K. O. Walden, Christopher Fields, John W. Bickham, J. Andrew Dewoody

## Abstract

Gray whales (*Eschrichtius robustus*) are unique as bottom feeding baleen whales and they have long been a conservation concern on both sides of the Pacific, in part because they migrate and disperse farther than any other species on earth. They experienced drastic population size declines due to environmental changes and commercial whaling. Here, we present an improved genome assembly for the gray whale. This genome assembly covers 2.4 Gb divided across 2689 contigs with an N50 of 15Mb. From the new assembly, we identify 75Mb sex-linked contigs and a identify 94.6% of searched genes based on Benchmarking Universal Single-Copy Ortholog score. We use the gray whale assembly to explore the effects of mapping to conspecific vs allospecific reference genomes when estimating genome-wide heterozygosity (*H*) and runs of homozygosity (ROH). The use of allospecific genomes significantly underestimate both *H* and ROH burden regardless of genomic distance and assembly quality. Our analyses highlight the importance of using contiguous conspecific assemblies in whale genomics and conservation.

**Significance:** Gray whales are unique in their behavior, morphology and ecology. The novel and contiguous genome assembly presented here will serve as a valuable resource for studies of their population and comparative genomics, as well as for identifying key adaptations that have evolved in this clade. Finally, our results demonstrate that biases can arise when using allopatric assemblies to evaluate diversity metrics, so they should be used and interpreted with caution.

## Introduction

High quality reference genome sequences provide excellent resources for the study of life on Earth (Blaxter, et al. 2022). Here we provide an improved genome assembly of the gray whale, which is the only extant baleen whale species adapted to bottom feeding and, because of its highly specialized morphology, is presently classified in the monotypic family Eschrichtidae. Gray whales are also remarkable in undertaking the longest annual migration (>20,000km) of any marine mammal during travel between Artic or Asian feeding grounds and eastern Pacific wintering grounds in Mexico (Mate, et al. 2015). Recently, a small number of transoceanic migrations from the Pacific to Atlantic Ocean basins (Hoelzel, et al. 2021) have been observed, as well as the first record of a gray whale in Hawaii (Baird et al., 2022), which illustrates the long-distance migratory capability of this species.

Reference genome assembly choice is important when addressing evolutionary and conservation genomic questions as the reference genome limits our ability to accurately estimate certain genomic metrics critical for reconstructing historical demography and estimating diversity. For many wildlife and non-model organisms a high-quality conspecific reference is simply not available and therefore mapping to either a low-quality conspecific or an allospecific reference genome are the only options. While a low quality conspecific genome assembly can provide robust estimates for some parameters (e.g., demographic changes in effective population size, Patton, et al. 2019), genome fragmentation makes it difficult to identify runs of homozygosity (ROH). The alternative is using an allospecific reference genome but this approach introduces bias in estimates in genome-wide heterozygosity (*H*), ROHs and demographic inference (Gopalakrishnan, et al. 2017; Prasad, et al. 2022).

In earlier gray whale studies, allospecific reference genomes have been used to quantify genomic diversity (Árnason, et al. 2018; Brüniche-Olsen, et al. 2018; Hoelzel, et al. 2021). Here, we present a PacBio HiFi gray whale genome and use this new gray whale genome assembly to explore how the choice of reference genome impacts genomic diversity estimates.

## Results and Discussion

### Genome assembly and annotation

We generated 474Gb of raw PacBio CLR read data and 149Gb of Illumina paired-end read data, using 90x final PacBio read coverage for the assembly, using all Illumina read data and PacBio data for polishing steps. The final assembly, after post-assembly polishing and filtering, covered 2.4Gb divided across 2689 contigs with an N50 of 15.0 Mb (Table 1), far more contiguous than our initial assembly (DeWoody, et al. 2017). We identified 38 contigs (75Mb) as being sex linked (Table S1), corresponding to ∼3.1% of the total assembly length. Benchmarking Universal Single-Copy Orthologs with the *laurasiatherian_odb10* database identified 94.6% of searched genes, substantially improved relative to earlier gray whale assemblies (DeWoody, et al. 2017).

**Table 1.**
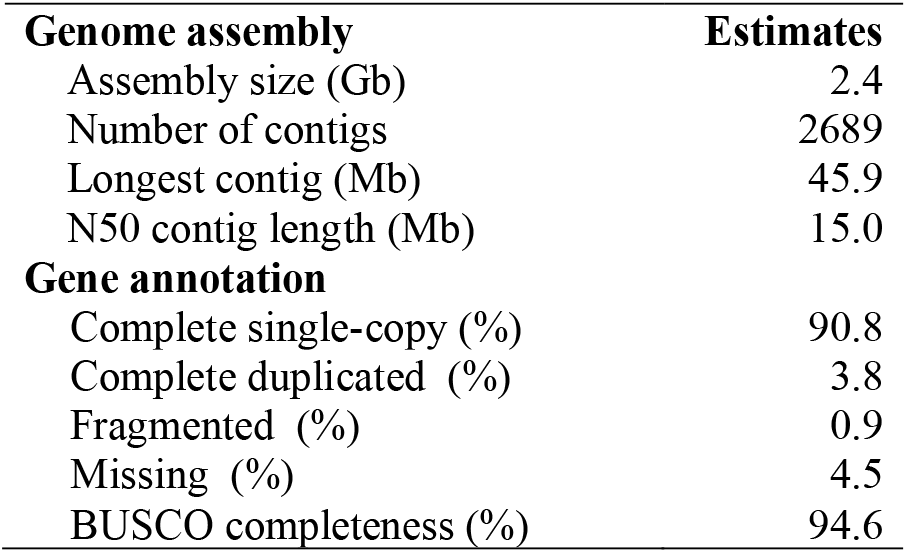
Genome assembly and annotation statistics for the gray whale.

### Heterozygosity and runs of homozygosity

Previous gray whale genomic studies have used an allospecific genome assembly as reference i.e., bowhead whale (*Balaena mysticetus*) (Árnason, et al. 2018), fin whale (*Balaenoptera physalus*) (Wolf, et al. 2022), and minke whale (*Balaenoptera acutorostrata*) (Brüniche-Olsen, et al. 2021) for mapping sequencing reads of gray whales. By mapping six gray whale genomes, our analyses show that the use of an allospecific reference genome underestimates mean individual *H* and ROH burden for the focal species independent of genomic distance to the allopatric refence (Figure 1 and Table S2). By mapping all six grey whale samples to our new gray whale genome asssembly, we estimated *H* to be 4.64×10^−4^ (±5.6×10^−6^; Figure 1a), which is higher than most baleens. Excluding regions with identified ROHs increased genome-wide *H* from 10-25% for all assemblies (Figure 1a). When we tested alternative estimates of *H* by allospecific mapping, however, we found consistently significantly lower *H* estimates, ranging from *H*= 3.7×10^−4^ (±2.0×10^−5^) when using the fin whale assembly to *H*= 4.0×10^−4^ (±3.6×10^−5^) when using the humpback whale assembly. These correspond to 15-20% lower *H* estimates when using an allospecific reference genome assembly compared to the gray whale genome assembly. This finding is consistent with results from *Canis spp*. where a ∼10% underestimation in *H* were identified when using an allopatric genome assembly (Gopalakrishnan, et al. 2017). The increase in bias we identify in baleen whales is likely due to the longer (∼7.49mya) divergence time (Table S2, Árnason, et al. 2018), compared to the relative recent (∼35kya) divergence time between dogs and wolves (Skoglund, et al. 2015). Interstingly increasing genomic distance to the focal species did not increase *H* estimates, as found in beluga whales (*Delphinapterus leucas*) (Prasad, et al. 2022).

**Figure 1.**
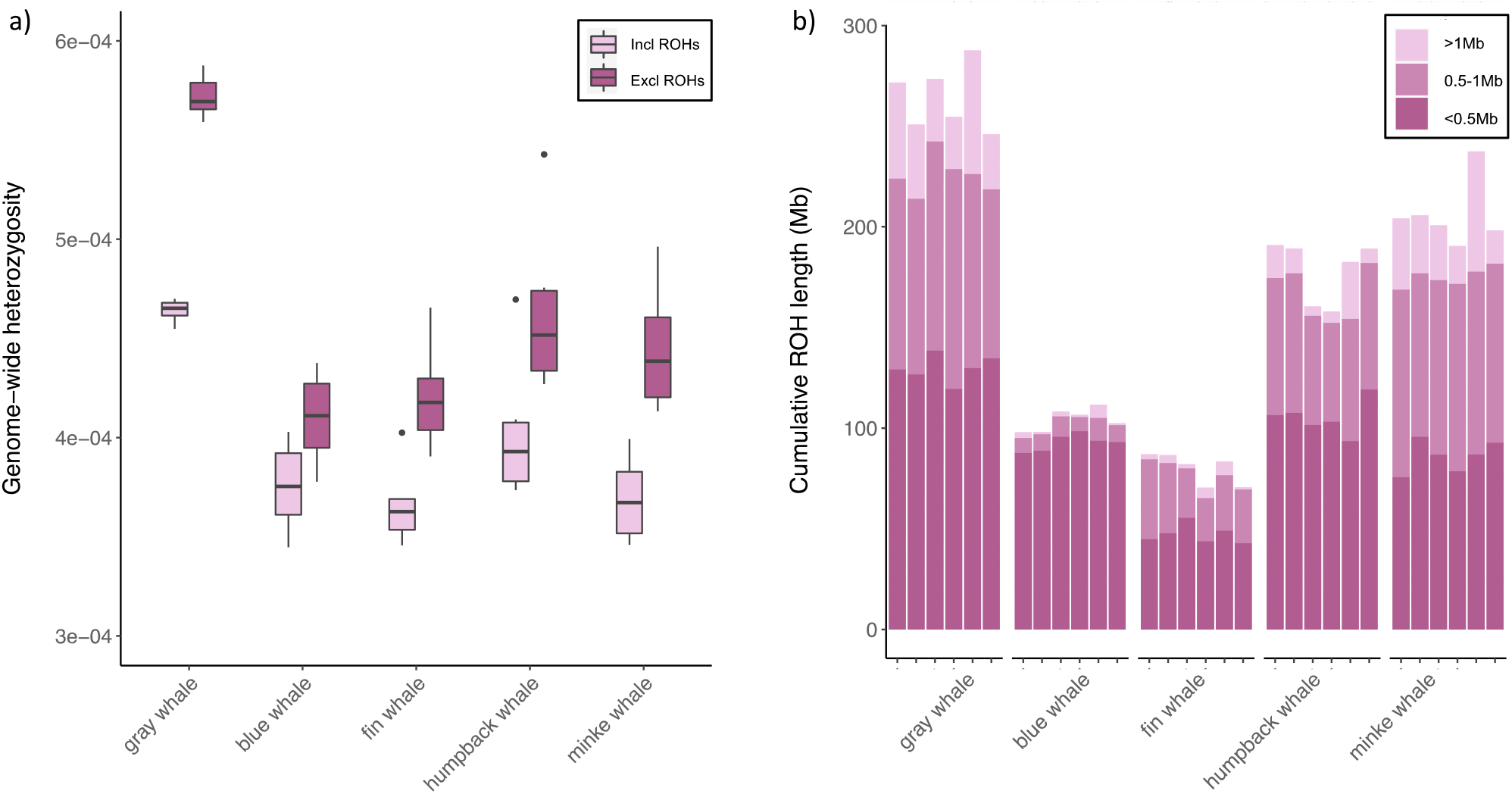
Mean heterozygosity and ROH burden for the six gray whales when mapped to a conspecific assembly and assemblies of increasing genomic distance. a) Genome-wide heterozygosity (*H*) estimates adjusted for inbreeding via the fraction of each individual’s genome in ROHs. *H* estimates including and excluding ROHs are shown side by side. Boxplots shows median, confidence intervals and outliers. b) ROH burden measured the fraction of ROHs per individual catogorised according to ROH length. Pairwise *t*-tests for the gray whale assembly to all other assemblies fo *H* and ROH burden, respectively, were significantly (*p* < 0.01) lower when mapping to an allospecific species.

ROHs are used in conservation genomics to assess the contribution of inbreeding to the overall genomic diversity and is a key measure for quantifying past population dynamics (Ceballos, et al. 2018). We normalized the amount of genome and number of sites surveyed to make the allospecific genomes comparable (Meyermans, et al. 2020). In all cases using an allospecific reference genome significantly underestimates the ROH burden (Figure 1b). Using a conspecific reference identified a higher number of ROHs for all classes (short <0.5Mb, medium <0.5-1Mb, and long >1Mb) and also contained a larger proportion of the genome in ROHs. We identify the smallest number of ROHs using the blue whale (*Balaenoptera musculus*) and fin whale reference genomes; in particular, the longer ROHs were not recovered using these allospecific references (Figure 1b). Blue and fin whale have similar genomic distances to the gray whale (Table S2), and represented assemblies in our dataset with the highest and lowest N50 respectively (Table S1), thus assembly N50 does not explain the difference in ROH pattern.

## Conclusion

We have generated an improved genome assembly for the gray whale. Our analyses show that the use of allospecific reference genomes underestimates genomic diversity estimates in the target species. The new gray whale assembly presented here will be an invaluable resource for future studies examining population and comparative genomics, particularly with regard to the evolution of unique genomic adaptations in this distinct clade (e.g., those related to benthic feeding behavior or long-distance movements).

## Materials and methods

### Genome sequencing, assembly and annotation

An eastern gray whale female (ER-17-0199) was chosen for constructing a *de novo* genome assembly (Brüniche-Olsen, et al. 2021). High molecular weight DNA was extracted by Polar Genomics and sequenced using Illumina and PacBio platforms. FASTQC v0.11.7 (Andrews 2017) was used throughout for quality assessment. All Illumina short-reads were processed with TRIMGALORE v0.6.5 (Krueger 2015) to remove adapters and low quality bases. PacBio subread binary alignment mapping files were converted over to FASTQ file format using SMRT LINK v8.0 (PacBio) command bam2fastq’ and concatenated. Reads <1kb or >50kb were removed prior to assembly using SEQKIT v0.12 (Shen, et al. 2016).

To generate a *de novo* assembly WTDBG2 v2.2 was used to create consensus contigs from a fuzzy *de Bruijn* graph (Ruan and Li 2020). PacBio reads were sub-sampled at different coverage levels (50X, 70x, and 90x) prior to assembly and QUAST v3.2 (Gurevich, et al. 2013) was used to assess quality (N50). PacBio subreads were mapped back to the optimal assembly with MINIMAP2 v2.11 (Li 2018), followed by consensus calling with ARROW v2.3 (Chin, et al. 2013). The Illumina short-reads were used for error correction using the POLCA script compiled with MASURCA v3.41 (Zimin, et al. 2017). To identify and remove highly heterozygous haplotypes assembled as separate primary contigs, we used the program PURGE_HAPLOTIGS v1.0 (Roach, et al. 2018). Primary haplotype sequences were then evaluated for contaminants using BLOBTOOLS2 V2.1 (Challis, et al. 2020). Genome assembly completeness was assessed with BUSCO v5.4.4 (Seppey, et al. 2019) comparing the assembly to the *laurasiatherian_odb10* database, containing 12,234 core orthologs. The assembly was annotated in three successive rounds with MAKER v 3.01.1 (Cantarel, et al. 2008) using: 1) published RNA-Seq datasets from liver and kidney tissue (Moskalev, et al. 2017; Toren, et al. 2020); 2) an alternative transcriptome assembly from beluga whale (NCBI TSA accession GGBT00000000); 3) protein predictions from a prior Illumina-only gray whale assembly as well as public fin whale, minke whale, and blue whale genome assemblies (NCBI accessions GCA_002738545.1, GCA_008795845.1, GCA_000493695.1, and GCA_009873245.2 respectively) [PMID: 29297306, 31553763], and 4) proteins from the curated SwissProt database (retrieved 2021-Jan) [PMID: 33237286]. Additional gene predictions were made using GENEMARK-EP+, part of the GENEMARK-ES v4.62 release (Brůna, et al. 2020) and ProtHint v2.5.0 using chordate protein sequences as input, and used as additional input in the final round of MAKER gene annotation.

### WGS mapping and genotyping

We downloaded whole genome resequencing SRAs from six published gray whale samples (Árnason, et al. 2018; Brüniche-Olsen, et al. 2018; DeWoody, et al. 2017; Hoelzel, et al. 2021). The cleaned reads were mapped to indexed genome assemblies in BWA v0.7.17 (Li and Durbin 2009) marking shorter splits as secondary (-M) and keeping individual read groups separate (-R). SAMTOOLS v1.9 (Li, et al. 2009) was used for indexing, sorting, fixing mates, marking duplicates, flagging reads that did not map correctly (view -F 3852). Genotypes were called with BCFTOOLS v1.15 (Danecek, et al. 2021; Li 2011) using the ‘mpileup’ command, omitting sites with low base (-Q 30) and mapping quality (-q 30) and using the multiallelic caller.

### Reference quality filtering

For each site in the reference genome, mappability was estimated using GENMAP (Pockrandt, et al. 2020). We used 100 bp k-mer allowing for two mismatches (-K 100 -E 2).

REPEATMASKER v4.1.2 (Smit, et al. 2015) was used to identify repeated segments (e.g., LINEs, SINEs, etc.) with the quick (-qq) option and ‘mammal’ repeat database (-species mammal). We identified sex-linked contigs in each of the genome assemblies using SATC (Nursyifa, et al. 2022). The idxstats input files were generated with SAMTOOLS. Sites with low mappability (mappability score <1), repeats, putative sex chromosome contigs, and contigs <1Mb were excluded from downstream analyses.

### Heterozygosity and runs of homozygosity

Genome-wide heterozygosity was estimated with ANGSD from the site frequency spectrum (- doCounts 1) using genotype likelihoods (-doMajorMinor 1). We used the GATK method (-GL 2) including only proper pairs (-only_proper_pairs 1), extended baq computation (-baq 2), removing low quality reads (-remove_bads 1 -C 50).) and filters on mapping and base quality (- minMapQ 30 -minQ 30). Only sites with a minimum of five reads were included in the SFS (- setMinDepthInd 5 -uniqueOnly 1).

ROHs were identified from the called genotypes with PLINK v1.90b6 (Purcell, et al. 2007). To compare ROHs across reference genomes, we thinned the VCF files accounting for differences in the number of sites passing MAF 0.01 filtering, sites passing reference genome QC, and thinning the number of sites according to the blue whale, which retained the smallest percentage of sites passing both reference genome QC and PLINK filtering (Table 3). We called ROHs using a minimum number of 20 SNPs (--homozyg-snp 20), minimum 10Kb in length (-- homozyg-kb 10), a SNP density of 1Mb (--homozyg-density 1000), and a window of 20 SNPs (-- homozyg-window-snp 20). We compared the means for each genomic diversity measure with paired *t*-tests in R v3.4.4 (Team 2019) (Figure 1).

### Genome-wide divergence

To estimate the divergence between the gray whales and the four baleen whale species (blue whale, humpback whale (*Megaptera novaeangliae*), minke whale, fin whale) we downloaded SRAs from Arnason *et al*. (2018) for each of the four species. The cleaned reads were mapped to the new gray whale genome assembly with BWA. ANGSD was used to estimated pairwise genetic distance (*d*_ij_) between individuals. We generated a consensus sequence for each individual (-doIBS 2) using filters on base quality (-minQ 30) and mapping (-minMapQ 30 - uniqueonly 1) while limiting the analyses to sites where all individuals had data (-minInd 4), discharging sites with low depth (-setMinDepthInd 5) and autosomal contigs >1Mb. The resulting distance matrix (-makeMatrix 1) was used to calculate genome-wide pairwise divergence between the baleen whales.

## Acknowledgements

ABO was supported by a Carlsberg Foundation Reintegration Fellowship (CF19-0427). JAD was supported in part by the U.S. National Institute of Food and Agriculture. Library construction and sequencing were conducted at the Roy J. Carver Biotechnology Center, University of Illinois at Urbana-Champaign. This work was supported in part by Exxon Neftegas Limited and Sakhalin Energy Investment Company. The funding parties had no part in study design, data collection, analysis or interpreting the results. The content herein is solely the responsibility of the authors and does not necessarily represent the official views of the funding parties.

## Data accessibility

The genome assembly has been uploaded to NCBI accession: NCBI WGS (accession JAIQCJ000000000, BioProject PRJNA707533). Previously published SRAs have accession numbers PRJNA384396, PRJNA389516 and PRJNA694958. The RNA-Seq datasets have accession number PRJNA706346.

