## Supplement material for "An improved gray whale assembly highlights how allospecific reference-genome choice can affect genomic diversity estimates"

### SUPPLEMENTARY MATERIAL

**Table S1.** Genome and mapping information for the genome assemblies.

| Species | contigs | N50 | Pre-filtering | Length >1Mb | Sites in repeats | Mappability <1 sites | Sex-linked contigs | Post-filtering | % kept |
|---|---|---|---|---|---|---|---|---|---|
| Gray whale | 2689 | 14870472 | 2431707408 | 2356318044 | 842747077 | 112664671 | 74742392 | 1400708387 | 58 |
| Blue whale | 130 | 110470125 | 2380012384 | 2374998319 | 1039025442 | 91256280 | 131757582 | 1235427446 | 52 |
| Fin whale | 62302 | 871016 | 2462774868 | 1080424691 | 611278757 | 102410 | 52205782 | 614904863 | 25 |
| Humpback whale | 2558 | 9138802 | 2265788366 | 2181271784 | 667476403 | 9535913 | 84932384 | 1353095872 | 60 |
| Minke whale | 10776 | 12843668 | 2431687698 | 2324429847 | 965496156 | 60266006 | 60051191 | 1215155055 | 50 |

**Table S2.** Genome-wide distance estimates between the gray whale and four rorquals. Short read sequences archived (SRAs) from each of the four species were mapped to the gray whale reference genome. Genome-wide distance was estimated on >1Mb autosomal contigs. * Divergence times are from Arnason *et al*. (2018).

| | Genetic distance | Divergence time (mya)* |
|---|---|---|
| Blue whale | 0.0130 | 8.35 |
| Fin whale | 0.0132 | 7.49 |
| Humpback whale | 0.0133 | 7.49 |
| Minke whale | 0.0149 | 10.48 |

**Table S3.** Site filtering for ROH calling. For each reference genome is listed the number of sites passing reference QC filtering, number of sites passing PLINK MAF>0.01 filtering, the percentage of sites that pass PLINK filtering, and the sites kept after thinning the number of sites comparable to the species with the smallest % of sites kept after filtering, in this case the blue whale genome which retained 0.011% of the reference QC sites after PLINK filtering.

| species | Sites passing reference QC filtering | Sites passing PLINK filtering | Percentage sites passing filtering | Sites kept after thinning |
|---|---|---|---|---|
| Gray whale | 1400708387 | 587991 | 0.042 | 153945 |
| Blue whale | 1235427446 | 135780 | 0.011 | 135780 |
| Fin whale | 614904863 | 269066 | 0.044 | 67581 |
| Humpback whale | 1353095872 | 665505 | 0.049 | 148712 |
| Minke whale | 1215155055 | 521708 | 0.043 | 133552 |
